# msiFlow 2.0: Extending Mass Spectrometry Imaging Workflows with Mass Recalibration, Annotation, Batch Correction, and Visual Quality Control

**DOI:** 10.64898/2026.11.11.765722

**Authors:** Philippa Spangenberg, Sebastian Bessler, Eva-Maria Hanschmann, Lars Widera, Jenny Bottek, Stephanie Thiebes, Alexander Potthoff, Olga Shevchuk, Sven Brandau, Jens Soltwisch, Daniel R. Engel

## Abstract

Mass spectrometry imaging (MSI) enables label-free, spatially resolved molecular profiling of tissues and cells, providing insights into physiological and pathological processes. However, its large and complex datasets require specialised computational approaches. We recently introduced msiFlow, an open source, end-to-end solution for reproducible and scalable MSI data analysis. Here, we present msiFlow 2.0, a major extension that integrates pixel-wise post-acquisition mass recalibration, automated lipid annotation, batch correction, and visual quality control, together with improved alignment, spatially informed deisotoping, and computational scalability. The workflow was evaluated using four human and mouse MALDI-2 and t-MALDI-2 MSI lipidomic datasets encompassing different biological conditions, sample types, and spatial resolutions. The updated computational mass recalibration and alignment workflow significantly improved mass accuracy of reported *m/z* values by correcting systematic mass deviations, achieving a mean absolute mass error below 1.5 ppm for independent MS/MS-supported lipid features. The integrated lipid annotation module reduced the number of unsupported candidate lipid assignments, while ComBat correction reduced sample-associated variation and retained separation of tissue compartments. Computational optimisation reduced alignment runtime by approximately 80%. Together, these advances provide an accurate, transparent, and scalable open-source framework for MSI data processing and downstream analysis.

**Graphical Summary:** 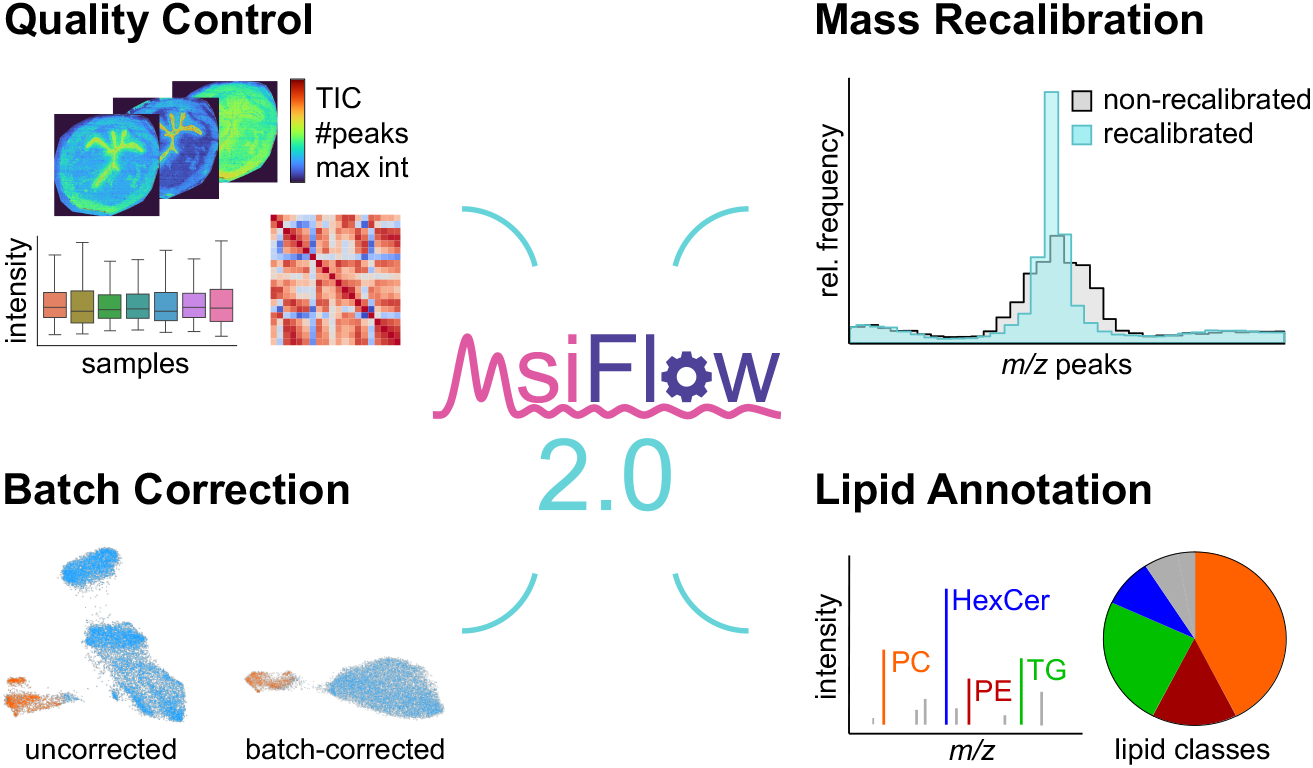

## Introduction

Matrix-assisted laser desorption/ionisation mass spectrometry imaging (MALDI MSI) enables label-free measurement of lipids and metabolites within their spatial context in tissues and cells. MALDI with laser-induced postionisation (MALDI-2) MSI^1^ further enhances molecular coverage, and transmission-mode (t-MALDI-2) MSI^2, 3^ additionally improves spatial resolution, providing deeper insights into the molecular landscape of physiological and pathological processes. However, extracting biological insights from raw MALDI MSI data remains challenging, as the resulting datasets are large, high-dimensional, and analytically complex, requiring specialised computational methods for data processing and downstream analysis. Furthermore, systematic variation across pixels and samples introduces batch-associated intensity differences that obscure biological patterns.

In MSI, a mass spectrum is acquired for each pixel across the tissue section, providing spatially resolved ion intensities that reflect relative analyte abundance as a function of mass-to-charge ratio (*m/z*). Comparative MSI data analysis and accurate molecular annotation require a consistent *m/z* vector across pixels and samples. However, experimental and instrumental variations can introduce systematic mass shifts, causing the same molecular ion to be detected at slightly different *m/z* values and complicating the construction of a common *m/z* vector. Reliable feature extraction further requires monoisotopic analyte peaks to be distinguished from isotope peaks and other signals arising from adducts, fragments, matrix-related ions, and background species. In addition, the increasing size and spatial resolution of MSI datasets place substantial demands on computational resources. Robust MSI preprocessing must therefore correct mass errors across pixels and samples, distinguish monoisotopic molecular features, account for batch-associated effects, and enable scalable data processing while preserving biologically meaningful variation^4^.

Consequently, numerous commercial and open-source software solutions have been developed for MSI data processing and analysis^5, 6^. Commercial platforms generally provide user-friendly graphical interfaces, but may offer limited customisability, transparency, and integration into automated and reproducible computational workflows. Open-source tools, such as BASIS^7^, M^2^aia^8^, HIT-MAP^9^, and massPix^10^, provide specialised functionality for preprocessing, multimodal image analysis, peptide identification, or lipid analysis. Other tools, including MSiReader^11^ and SpectralAnalysis^12^, depend on proprietary software environments, such as MATLAB, whereas comprehensive frameworks such as Cardinal^13, 14^ provide broader functionality but require programming expertise. Consequently, complete MSI analyses frequently rely on combinations of commercial platforms, open-source packages, and in-house scripts, complicating reproducible and high-throughput application and creating barriers for non-expert users. Workflow-based approaches, including Galaxy^15, 16^ and Snakemake^17^, aim to facilitate the integration of diverse MSI analysis methods into reproducible and user-defined workflows.

Available MSI preprocessing tools commonly include normalisation and spectral alignment. Normalisation can reduce pixel- and sample-associated variation in signal intensity but may not adequately account for systematic differences between experimental batches^7, 18^. Recent comparative studies identified ComBat^19^ as a readily applicable method for reducing batch-related variance in MALDI MSI while retaining biologically meaningful variation under appropriate experimental conditions^20, 21^. Alignment enables corresponding features across spectra to be mapped onto a common *m/z* vector but does not necessarily correct systematic deviations from their theoretical masses. Complementary mass recalibration using matrix-derived or endogenous reference ions can compensate for such deviations, improve mass accuracy, and thereby increase the reliability of molecular annotation^22, 23^. To integrate MSI processing and downstream analysis within a reproducible framework, we previously introduced msiFlow 1.0^24^, an open-source, end-to-end software solution for MALDI MSI data analysis combining preprocessing, registration, statistical analysis, and visualisation in automated and scalable workflows. Its configuration-driven operation does not require specialised programming expertise, although basic familiarity with containerised software environments such as Docker^25^ is needed for deployment and execution.

Although msiFlow 1.0 transforms raw MSI data into standardised spectra suitable for downstream analysis, several limitations remained. First, msiFlow 1.0 uses an unsupervised peak frequency-based approach to construct a common *m/z* vector across all pixel spectra. Although this represents an established strategy for MSI alignment, generating the global peak-frequency distribution in msiFlow 1.0 requires all detected *m/z* values to be combined into a single in-memory array, resulting in high memory requirements and increasing computational costs for large datasets. Moreover, msiFlow 1.0 does not include a dedicated post-acquisition mass recalibration step to correct systematic deviations from theoretical reference masses before alignment. Second, the deisotoping algorithm implemented in msiFlow 1.0 identifies isotope peaks based primarily on their expected mass differences and relative intensities. It does not account for the similarity in spatial intensity distribution of monoisotopic and isotope peaks across pixels, although their spatial distributions can provide complementary evidence for isotope assignment. Third, lipid annotation is not directly integrated into the msiFlow 1.0 workflow. Monoisotopic peak lists must instead be submitted to the external LIPID MAPS bulk structure search, after which the resulting annotation table can optionally be reintroduced into msiFlow 1.0 for downstream analysis. This external step interrupts the automated workflow and provides limited control over workflow-specific database generation and mass-matching criteria. Finally, while msiFlow 1.0 includes normalisation methods to reduce pixel- and sample-associated intensity variation, it does not provide a dedicated method for correcting systematic batch effects.

To address these limitations, we present msiFlow 2.0, a major update that extends the previous version by improving analytical accuracy, computational scalability, efficiency and transparency. msiFlow 2.0 integrates new modules for post-acquisition lock-mass-based recalibration, automated lipid annotation, and batch correction, together with improved alignment and deisotoping procedures. Memory-efficient and parallelised processing reduces the computational requirements of large MSI datasets. Evaluation across four human and mouse MALDI-2 and t-MALDI-2 MSI lipidomic datasets demonstrates the applicability of the extended workflow to different disease states, sample types, and spatial resolutions.

## Experimental Section

### Patient Ethics

Tumour specimens were collected during surgical resection from patients with head and neck squamous cell carcinoma (HNSCC), in accordance with the Declaration of Helsinki and following approval by the local ethics committee at the medical faculty of the University Duisburg-Essen. Written informed consent was obtained from all patients prior to sample collection. Patient characteristics are provided in Table S1.

### Sample Preparation

Tumour samples obtained from HNSCC patients were immediately stored in Ringer’s solution and processed within the next hour. Processing involved embedding the samples in Tissue-Tek O.C.T. Compound (Sakura Finetek, Umkirchen, Germany) and freezing them using liquid nitrogen. Tissue sections with a thickness of 5 µm were prepared using a cryostat. Samples were stored at -80°C until use.

Sample preparation of mouse bladders was described in detail previously^26^. Briefly, bladders were harvested 24 h post-infection with uropathogenic *Escherichia coli* (UPEC; strain 536 (O6:K15:H31)) from mice that received either UPEC infection alone or additional TSP1/TGF-β1 inhibition 1 h post-infection. The harvested tissues were immediately snap-frozen in 2-methylbutane and cryosectioned at 8 µm thickness. Sections were mounted onto SuperFrost™ slides, vacuum-sealed and stored at −80 °C until further processing.

For immunofluorescence staining, sections were equilibrated to room temperature for 10 min, after which all subsequent steps were performed at 4 °C. Sections were fixed in 4% formalin for 5 min and washed once with cold PBS for 30 s. An antibody solution prepared in 1% BSA in PBS was applied to each section and incubated for 1 h, followed by three washing steps with PBS for 30 s. For nuclear stain, sections were incubated with DAPI (1:5,000; Life Technologies, Cat. No. D1306), diluted in 1% BSA in PBS for 5 min. Finally, sections were washed three times with 150 mM ammonium acetate for 30 s. Samples were maintained on dry ice during transportation.

### MALDI-2 Mass Spectrometry Imaging

HNSCC tissue sections were thawed under a gentle stream of dry N₂ gas. To remove the OCT embedding medium, the tissue sections were dipped in 150 mM ammonium formate solution at 4°C according to the protocol described by Truong et al.^27^. The sections were then dried under a gentle stream of dry N₂ gas.

The 2,5-DHAP matrix was applied to the tissue sections by sublimation using a home-built sublimation device described previously by Potthoff et al.^28^. Briefly, 0.75 mL of a 20 mg/mL solution of 2,5-DHAP in acetone was introduced into the matrix reservoir and heated to approximately 100 °C to evaporate the acetone. The sample was then clamped onto the cold side of the device at room temperature, and the entire system was sealed and evacuated to around 10^-1^ mbar. The cold side was first cooled to approximately −15 °C, and then the pressure was set below 10^-3^ mbar to initiate matrix sublimation and resublimation onto the tissue section. After 10 min, heating of the hot side was discontinued, and the cold side was warmed to room temperature before the system was vented to prevent condensation of ambient moisture on the sample. The samples were measured immediately after matrix preparation.

MALDI-2 MSI data were acquired at a pixel size of 10 µm using a timsTOF fleX MALDI-2 instrument equipped with the microGRID extension (Bruker Daltonics, Bremen, Germany). All MALDI MSI measurements were performed in positive ion mode with TIMS disabled. The MALDI and postionisation lasers were operated at a pulse repetition rate of 1 kHz with a delay of 10 µs. The ablation laser power was set to 70%, with 24 shots acquired per pixel and beam scan enabled and set to 9x9 µm^2^. The ion detection range was set to *m/z* 300–1500. Mass recalibration was performed before each measurement using red phosphorus.

### Transmission-mode MALDI-2 Mass Spectrometry Imaging

High-resolution t-MALDI-2 MSI data of mouse bladder tissue sections were acquired using a prototype transmission-mode MALDI ion source coupled to a timsTOF fleX MALDI-2 mass spectrometer (Bruker Daltonics, Bremen, Germany), as described previously by Potthoff et al.^28^ The instrument was operated in positive ion mode with trapped ion mobility spectrometry (TIMS) disabled. MALDI and postionisation lasers were operated at a pulse repetition rate of 1 kHz with an inter-laser delay of 10 µs. Mass spectra were acquired using 25 laser shots per pixel over an *m/z* range of 300–1500. Imaging data were recorded at a pixel size of 2 × 2 µm².

### Datasets

msiFlow 2.0 was developed and evaluated using four spatial lipidomic datasets acquired from human and mouse samples by MALDI-2 and t-MALDI-2 MSI. The datasets included whole-tissue and single-cell measurements, pixel sizes ranging from 2 to 10 µm, and different biological and disease conditions. The first dataset comprised head and neck cancer (HNC) tissue samples (*n* = 29, 882 GB). The second, previously published dataset comprised single-cell MSI data from blood-derived neutrophils obtained from healthy individuals and patients with liver cirrhosis (three healthy controls and three patients with liver cirrhosis, *n* = 6 biological samples across three measurements, 20 GB)^29^. The third dataset comprised mouse bladder tissue sections from four experimental conditions: uninfected control, uropathogenic *E. coli* (UPEC)-infection, and the corresponding groups treated with transforming growth factor beta 1 (TGF-β1) inhibitor (*n* = 20, 347 GB). These samples were acquired by MALDI-2 MSI at a pixel size of 5 µm^26^. Finally, a high-resolution subset of the mouse bladder samples prepared for multimodal imaging was additionally acquired by t-MALDI-2 MSI with integrated in-source microscopy at a pixel size of 2 µm. Only the MSI data were included in the present study and analysed as a fourth dataset (*n* = 2, 313 GB)^26^.

### Data Preprocessing

All datasets were processed using msiFlow 1.0 and msiFlow 2.0 on AMD Zen 4-based computing nodes of the PALMA II high-performance computing cluster at the University of Münster. Each node was equipped with 192 CPU cores and 722 GB of memory. The software versions, workflow-specific parameters, and computational resource settings used for each dataset are provided in the configuration files deposited on GitHub. The preprocessing workflow implemented in msiFlow 1.0 has been described in detail previously^24^. Therefore, the following sections focus on the newly implemented and updated processing steps introduced in msiFlow 2.0.

### Software Implementation

msiFlow 2.0 represents the next major release of msiFlow, incorporating new functionality, expanded preprocessing capabilities, and performance improvements while preserving the modular design of the original software. Detailed descriptions of the msiFlow 1.0 software design and functionality have been reported previously^24^. Briefly, msiFlow was implemented in Python and developed as a collection of computational workflows for MSI data preprocessing and analysis. The workflows were automated and standardised using Snakemake^17^, a Python-based workflow management system that enables reproducible and scalable data analysis by defining individual processing steps as rules with specified inputs, outputs, and execution commands. In msiFlow, execution commands invoke a dedicated Python script through the command-line interface. Each script is designed to perform a single processing step and accept user-defined parameters, which are specified in a central configuration file. msiFlow 2.0 represents a major update of msiFlow by introducing new preprocessing steps, including post-acquisition mass recalibration and annotation, while enhancing the functionality of the existing alignment and deisotoping procedures. Within the segmentation workflow, a batch correction step was added to account for batch-associated variations. In addition, computational performance of the alignment step was improved, and new modules were implemented for the generation of multiple visualisation outputs to support quality control and data assessment. The preprocessing workflow accepts raw MSI data (in d and imzML format) and produces standardised imzML files, monoisotopic feature lists with candidate lipid annotations, OME-TIFF image stacks, and quality-control outputs for downstream analysis. The source code, software environment, and configuration files corresponding to the analyses presented in this study are publicly available, as detailed in the Code Availability Statement.

### Post-acquisition Mass Recalibration

msiFlow 2.0 introduces a pixel-wise mass-recalibration step into the preprocessing workflow to correct the average mass error arising from experimental and technical variation. For each pixel spectrum, a recalibration function is calculated from the deviations between selected experimental peaks and their corresponding predefined reference masses. Depending on the selected method, mass errors are corrected by either median correction or linear regression. Experimental peaks are matched to the reference masses based on three selection criteria: (1) peak has the highest intensity within a user-defined mass window, (2) peak exceeds a user-defined minimum intensity threshold, (3) peak lies within a user-defined maximum mass shift from the reference mass specified in ppm or Dalton. Reference masses at *m/z* 369.351578 (Cholesterol, C_27_H_45_O^+^, [M-H_2_O+H]^+^), 577.519037 (DG 34:1, C_37_H_69_O_4_^+^, [M-H_2_O+H]^+^), and 760.585082 (PC 34:1, C_42_H_83_NO_8_P^+^, [M+H]^+^) were used for all four datasets. The cholesterol-derived ion was selected as reference, as cholesterol is normally contained in all membranes, and its ion exhibits high signal intensity in the analysed datasets. PC 34:1 is a commonly detected lipid that exhibits high signal intensity in the analysed datasets, making it suitable as a reference. During the MALDI and MALDI-2 process *m/z* 577.519037 is produced from 34:1 lipids (mostly as a fragment of PC 34:1) showing comparable spatial distribution to PC 34:1, making it suitable as additional reference. All three reference masses were validated by MS/MS in a previous study^29^. A user-defined parameter specifies the minimum number of reference masses that must be detected for a pixel to be recalibrated. In the present study, this threshold was set to the total number of reference masses. Consequently, recalibration was performed only for pixel spectra in which the peak-selection criteria were fulfilled for all predefined reference masses. Pixel spectra that did not meet this requirement were not recalibrated and are indicated in grey in the quality-control outputs. To assess recalibration performance, msiFlow 2.0 generates spatial heatmaps showing the median mass error across all reference masses for each pixel before and after recalibration. These heatmaps additionally identify pixels that do not fulfil the recalibration criteria. Visual assessment of the resulting maps showed that most on-tissue pixels were successfully recalibrated across all four datasets. Manual inspection of representative raw pixel spectra revealed mass deviations of up to 30 ppm. Accordingly, 30 ppm was selected as the maximum allowable deviation for matching experimental peaks to the predefined reference masses.

### Alignment

Pixel-wise mass recalibration was performed before construction of the common *m/z* vector and spectral alignment. The alignment step of the preprocessing workflow was updated in msiFlow 2.0 to improve the construction of the common *m/z* vector and computational performance. Whereas recalibration corrects technically and experimentally induced mass shifts within individual pixel spectra relative to predefined reference masses, alignment maps corresponding peaks across all pixels of a dataset to a common *m/z* vector. To construct this, an adapted kernel-based density approach based on BASIS^7^ is used. First, a histogram of all detected *m/z* values is generated using a user-defined bin width. The resulting peak-frequency distribution is then smoothed using locally weighted scatterplot smoothing (LOESS), and the local maxima of the smoothed distribution are selected to form the common *m/z* values. The measured peaks are subsequently mapped to the common *m/z* vector using the nearest-neighbour matching approach implemented in BASIS. A user-defined mass tolerance restricts the mapping to peaks within the specified distance from a common *m/z* value. While the histogram bin size and the mass tolerance were specified in Dalton in msiFlow 1.0, the updated version of msiFlow 2.0 enables these parameters to be defined in ppm, thereby accounting for the *m/z*-dependent absolute width of a constant relative mass tolerance. To reduce memory requirements and runtime, msiFlow 2.0 introduces a chunked data processing mode. In the standard in-memory mode, the detected peaks from all selected pixel spectra are combined into a single array to generate the histogram of peak frequencies. In the chunked processing mode, the global *m/z* array is partitioned into smaller data chunks containing a predefined number of pixels. For each data chunk, a histogram is computed and the resulting histograms are subsequently aggregated to obtain the final histogram across all pixels. In addition, Python’s multiprocessing module is used for parallelising data import in the default processing mode and for parallelising histogram computation across data chunks in the chunked processing mode. Hence, the minimum memory requirement in chunked processing mode depends on the selected chunk size, which is dependent on the number of peaks per pixel spectrum, rather than on the total dataset size. The number of pixels per chunk and the corresponding memory requirement can be specified in the configuration, together with the available CPU cores and system memory. Based on these settings, the workflow automatically selects the number of processes that can be run in parallel.

### Deisotoping

In msiFlow 1.0, isotope peaks were identified based on their expected mass differences and relative intensities. A candidate peak was classified as an isotope if its deviation from the expected isotope position was within a user-defined mass tolerance and its intensity was lower than that of the preceding peak in the putative isotope series. In msiFlow 2.0, the deisotoping algorithm was extended by incorporating spatial similarities as an additional criterion for isotope assignment. The updated algorithm employs an iterative search strategy to identify consecutive peaks within a putative isotope series. At each iteration, a candidate peak is evaluated according to three criteria: (1) its deviation from the expected isotope *m/z* value is within a user-defined mass tolerance, (2) its spatial intensity distribution is sufficiently similar to that of the putative monoisotopic peak, as determined by Pearson’s correlation coefficient, (3) its intensity is lower than that of the preceding peak but exceeds a user-defined minimum fraction of the preceding peak intensity. In the present study, the mass tolerance was set to 5 ppm, corresponding to the expected mass accuracy and the minimum Pearson correlation coefficient was set to 0.4 to require a degree of similarity between the spatial intensity distributions of candidate isotope peaks and the putative monoisotopic peak. The minimum relative intensity was set to 20% of the preceding peak intensity, reflecting the generally decreasing pattern of consecutive isotope peaks. If all criteria are fulfilled, the candidate is classified as an isotope peak, and the search is continued iteratively to identify subsequent isotope peaks. For quality control, msiFlow 2.0 creates a table containing all isotope candidates with their three measured criteria.

### Lipid Annotation

msiFlow 2.0 introduces an automated, accurate-mass-based MS1 lipid annotation step to the preprocessing workflow. A user-defined lipid reference database is generated from a structure-data file obtained from the LIPID MAPS Structure Database (https://www.lipidmaps.org/databases/lmsd/download). The file is parsed using the open-source cheminformatics toolkit RDKit (https://www.rdkit.org). Lipid entries are filtered according to user-defined lipid classes and adduct types, and relevant information, including lipid name, lipid class, molecular formula, exact neutral mass, adduct type, and calculated adduct *m/z*, is extracted. Following database generation, the measured *m/z* values are matched to the calculated adduct *m/z* values using a user-defined mass tolerance specified in ppm or Dalton. Additionally, the search can be restricted to all lipid species with an even total carbon number or performed without carbon-number filtering. All candidate matches are reported in an annotation table containing the measured *m/z*, lipid name, lipid class, molecular formula, adduct type, exact neutral mass, calculated adduct *m/z*, and corresponding mass error in ppm or Dalton. Because the annotation is based on MS1 accurate-mass matching alone, multiple candidate lipids may be assigned to the same measured *m/z* value.

### Batch Correction

To reduce systematic intensity variation associated with different experimental batches, a batch correction step based on the Python implementation of ComBat^30^ was integrated into the segmentation workflow of msiFlow 2.0. For the analysis presented here, the individual sample identifiers were specified as batch labels because pixels originating from the same tissue section share sample-specific technical effects related to sample preparation, acquisition, and signal intensity. ComBat estimates feature-specific batch effects using an empirical Bayes framework and adjusts the feature intensities accordingly. Batch correction was performed before dimensionality reduction and clustering. Where applicable, biological variation can be included in the ComBat model to reduce the risk of removing variation associated with the experimental conditions. In this study, biological covariates were not included in the model. Therefore, the correction was intended to reduce sample-associated variation but may also affect biological differences that are confounded with sample identity.

### Statistical Analysis

Differences in mass error of reported *m/z* values between msiFlow 1.0 and msiFlow 2.0 versions (updated and new) were statistically analysed in GraphPad Prism 11. Only *m/z* features detected by all three software versions were included in the analysis. Absolute mass errors were analysed using a Friedman test for repeated measures, followed by Dunn’s multiple comparisons test to evaluate pairwise differences between software versions. Data are presented as median with interquartile range (IQR), and adjusted p-values < 0.05 were considered statistically significant. To assess computational performance of msiFlow 2.0 compared to msiFlow 1.0, the bladder and HNC dataset were processed five times independently using each version. Runtime and memory usage of msiFlow 1.0 and msiFlow 2.0 were recorded for each processing run and statistically compared using a two-tailed Mann-Whitney test. A p-value < 0.05 was considered statistically significant.

### Neutrophil Clustering of Bladder Data

msiFlow 1.0 was used to analyse the spatial lipidome of neutrophils in urinary tract infection in a recent study^26^. As part of this study, neutrophil-associated pixels were identified and extracted from the MALDI-2 MSI dataset. The extracted pixel spectra and their assignments to the urothelium or connective and muscle tissue were used here to evaluate the batch-correction module implemented in msiFlow 2.0. The neutrophil-associated MSI data were processed using the segmentation workflows of msiFlow 1.0 and msiFlow 2.0. In both workflows, dimensionality reduction was performed using uniform manifold approximation and projection (UMAP)^31^, followed by Hierarchical Density-Based Spatial Clustering of Applications with Noise (HDBSCAN)^32^. For the msiFlow 2.0 analysis, ComBat batch correction was applied to the feature-intensity matrix before UMAP dimensionality reduction, using the individual sample identifiers as batch labels. Tissue-compartment assignments were not used for batch correction but were retained for subsequent evaluation of the UMAP embeddings. The msiFlow 1.0 analysis was performed without batch correction. All other preprocessing, UMAP, and HDBSCAN parameters were kept identical between the two analyses.

## Results and Discussion

### Overview of the msiFlow 2.0 Workflow

msiFlow 2.0 extends the original workflow by introducing lock-mass recalibration and automated lipid annotation, while improving spectral alignment, deisotoping, and computational performance (Figure 1A). Together, these developments enable the generation of standardised MSI datasets containing monoisotopic molecular features with improved mass accuracy and associated candidate lipid annotations. In addition, msiFlow 2.0 integrates batch correction and multiple visualisation outputs for transparent and efficient data processing as well as quality assessment.

**Figure 1.**
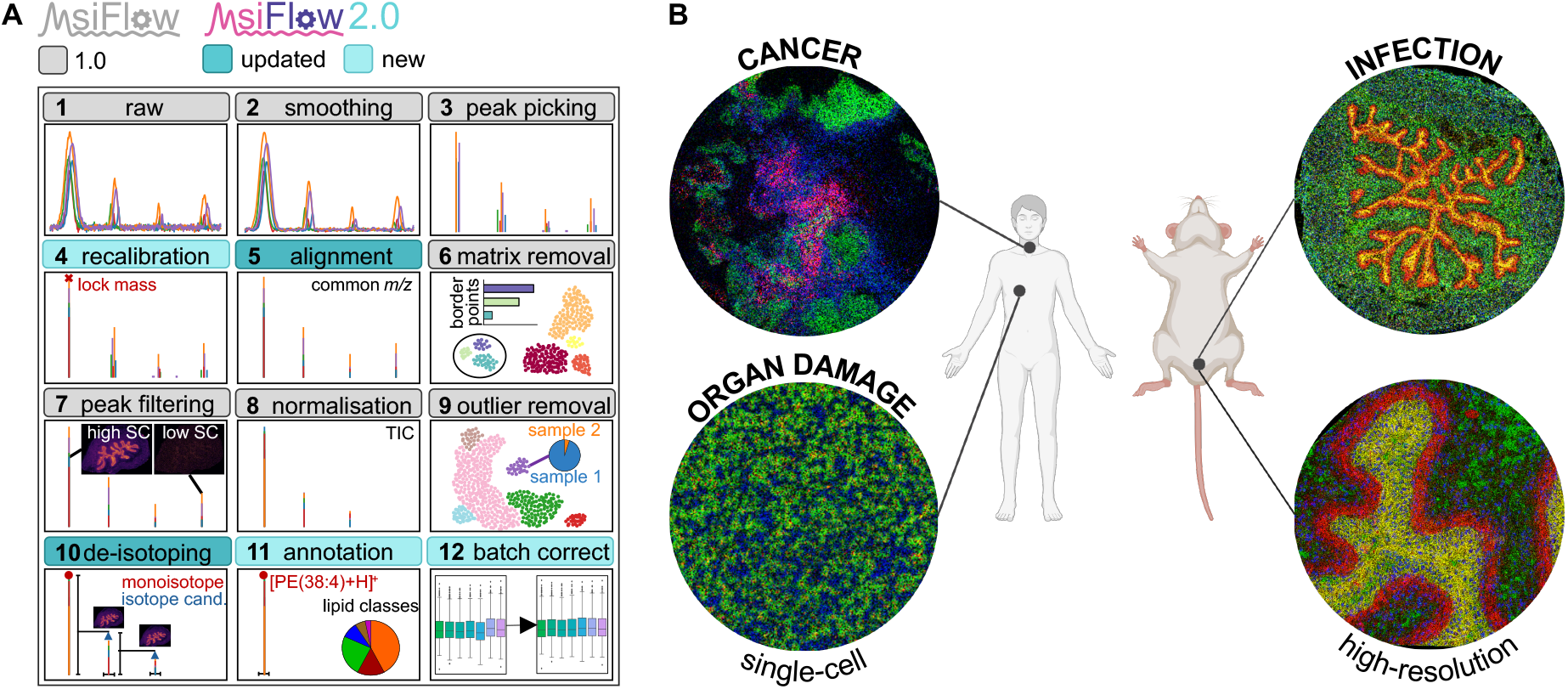
Overview of the preprocessing modules in msiFlow 2.0 and the datasets used for its evaluation. (**A**) Schematic overview of the msiFlow 2.0 modules of the preprocessing workflow (1-11) and batch correction (12), indicating modules retained from msiFlow 1.0 (light grey), enhanced modules (dark turquoise), and newly implemented modules (light turquoise). (**B**) Overview of the four MSI datasets used to evaluate msiFlow 2.0. Human head and neck cancer tissue (top left; *n* = 29), blood-derived neutrophils obtained from healthy individuals and patients with liver cirrhosis (bottom left; *n* = 6), mouse bladder tissue from urinary tract infection analysed by MALDI-2 MSI (top right; *n* = 20), and a high-resolution mouse bladder dataset acquired by t-MALDI-2 MSI (bottom right; *n* = 2). SC, spatial coherence; TIC, total ion current; cand., candidate. Figure partially created in BioRender. Engel, D. (2026) https://BioRender.com/3yoc6l8.

To improve mass accuracy, msiFlow 2.0 introduces pixel-wise recalibration based on user-defined reference masses (Figure 1A, step 4) and an updated alignment procedure (Figure 1A, step 5). Recalibration corrects systematic mass deviations within individual pixel spectra, whereas alignment subsequently maps corresponding peaks across spectra onto a common *m/z* vector. In msiFlow 2.0, this common vector is derived from the local maxima of a smoothed peak-frequency distribution. The updated alignment supports ppm-based bin widths and matching tolerances, which better reflect the relative mass accuracy of high-resolution mass spectrometers across the analysed mass range than fixed tolerances in Dalton. Together, recalibration and ppm-based alignment are expected to reduce artificial feature splitting and improve the consistency of corresponding molecular features across pixels and samples.

msiFlow 2.0 further extends deisotoping and integrates automated lipid annotation. The updated deisotoping algorithm (Figure 1A, step 10) combines expected isotope mass differences and relative intensities with the spatial similarity between candidate isotope and monoisotopic peaks. As isotopic ions originating from the same molecular species are expected to exhibit similar spatial distributions, spatial correlation provides complementary evidence for isotope assignment and may reduce classifications based solely on coincidental mass and intensity relationships. This concept is related to metabolite-annotation approaches such as METASPACE^33^, which combines mass accuracy, isotope patterns, and spatial distribution as well as related approaches based on mass and intensity information^10, 34^. However, the current algorithm assumes a decreasing isotope-intensity pattern and its performance remains dependent on the selected mass, intensity, and correlation thresholds. Unlike approaches that use molecular formulas to predict isotope distributions, the present method enables annotation-independent deisotoping for untargeted MSI data preprocessing without requiring prior knowledge of molecular identity or elemental composition.The automated annotation module (Figure 1A, step 11) generates a lipid reference library and matches measured *m/z* values to calculated adduct *m/z* values according to user-defined criteria, including mass tolerance, lipid class, adduct type, and carbon-number filtering. Similar to the annotation approach of massPix^10^, this algorithm facilitates systematic and reproducible annotation across datasets. Its integration into msiFlow 2.0 avoids the external database-search and data-reimport steps required in msiFlow 1.0. Nevertheless, because the assignments are based primarily on accurate-mass matching, they represent candidate lipid annotations and cannot distinguish isobaric or isomeric lipid structures without orthogonal evidence such as MS/MS. In addition, while the optional carbon-number filter can reduce the number of candidate annotations, it may exclude biologically relevant odd-chain lipid species if present in the measured data. Therefore, the filter should be selected according to the expected lipid composition and research question, considering which lipid species are detectable in the dataset.

To minimise systematic intensity variations introduced by different experimental runs, msiFlow 2.0 integrates the Python implementation of ComBat^30^ as batch correction method (Figure 1A, step 12). Comparative studies have identified ComBat as a readily applicable batch-correction method for MALDI MSI^20, 21^. Its integration into msiFlow enables batch-associated variation to be addressed within the same reproducible workflow. However, successful batch correction requires an experimental design in which technical batch and biological condition are not fully confounded, as otherwise relevant biological variation may also be removed.

Visual quality-control outputs were implemented throughout the workflow to facilitate assessment of both data quality and processing parameters. Spatial and spectral visualisations support assessment of data quality, smoothing, peak picking, recalibration, and alignment (Figures S1-S3). These outputs are particularly relevant for parameter-dependent processing steps, as they enable users to identify inappropriate settings or local processing failures rather than relying solely on final summary measures. The final preprocessing outputs comprise standardised imzML files, a list of monoisotopic molecular features with candidate lipid annotations, and OME-TIFF image stacks of analyte distributions. In addition, msiFlow 2.0 generates sample-level PCA plots, heatmaps, and box plots of summarised feature intensities, together with summaries of annotated lipid classes and candidate annotations per measured *m/z* (Figure S4). Overall, the visual quality-control outputs enable users to continuously assess data quality at multiple stages of the workflow and identify potential issues arising from preprocessing or inappropriate parameter settings.

In addition to the analytical and functional extensions, msiFlow 2.0 includes computational optimisations designed to address the increasing size and computational demands of high-resolution MSI datasets. In msiFlow 1.0, generating the common *m/z* vector required loading all peaks into one large in-memory array, resulting in high memory requirements and a substantial computational bottleneck. In msiFlow 2.0, the data is processed in parallel in smaller data pieces that fit into available memory. These changes improve computational scalability and facilitate the analysis of datasets containing large numbers of pixels and detected peaks. Their effect on runtime was evaluated quantitatively in the subsequent performance analysis.

To evaluate msiFlow 2.0 across different MSI settings, we applied the extended workflow to four human and mouse spatial lipidomic datasets acquired by MALDI-2 and t-MALDI-2 MSI (Figure 1B). These comprised human head and neck cancer tissue (*n* = 29), blood-derived neutrophils from healthy individuals and patients with liver cirrhosis (*n* = 6), mouse bladder tissue from urinary tract infection experiments analysed by MALDI-2 MSI (*n* = 20), and a high-resolution mouse bladder dataset acquired by t-MALDI-2 MSI (*n* = 2). Thus, the evaluation included whole-tissue and single-cell measurements, different biological conditions, and spatial resolutions ranging from 2 to 10 µm.

### Mass Recalibration Improves Accuracy of the Common *m/z* Vector

msiFlow 2.0 introduces a pixel-wise mass recalibration step to correct systematic mass deviations arising from instrumental and experimental variation. Similar to established approaches based on matrix-derived or endogenous reference ions^22, 23^, the algorithm matches experimental peaks to predefined reference masses and uses their mass deviations to calculate a pixel-specific recalibration function based on three selection criteria (Figure 2A). Candidate reference peaks are selected according to their position within a user-defined mass window, minimum intensity, and maximum allowable mass deviation. Depending on the selected method, either median correction or linear regression is subsequently applied to the respective pixel spectrum.

**Figure 2.**
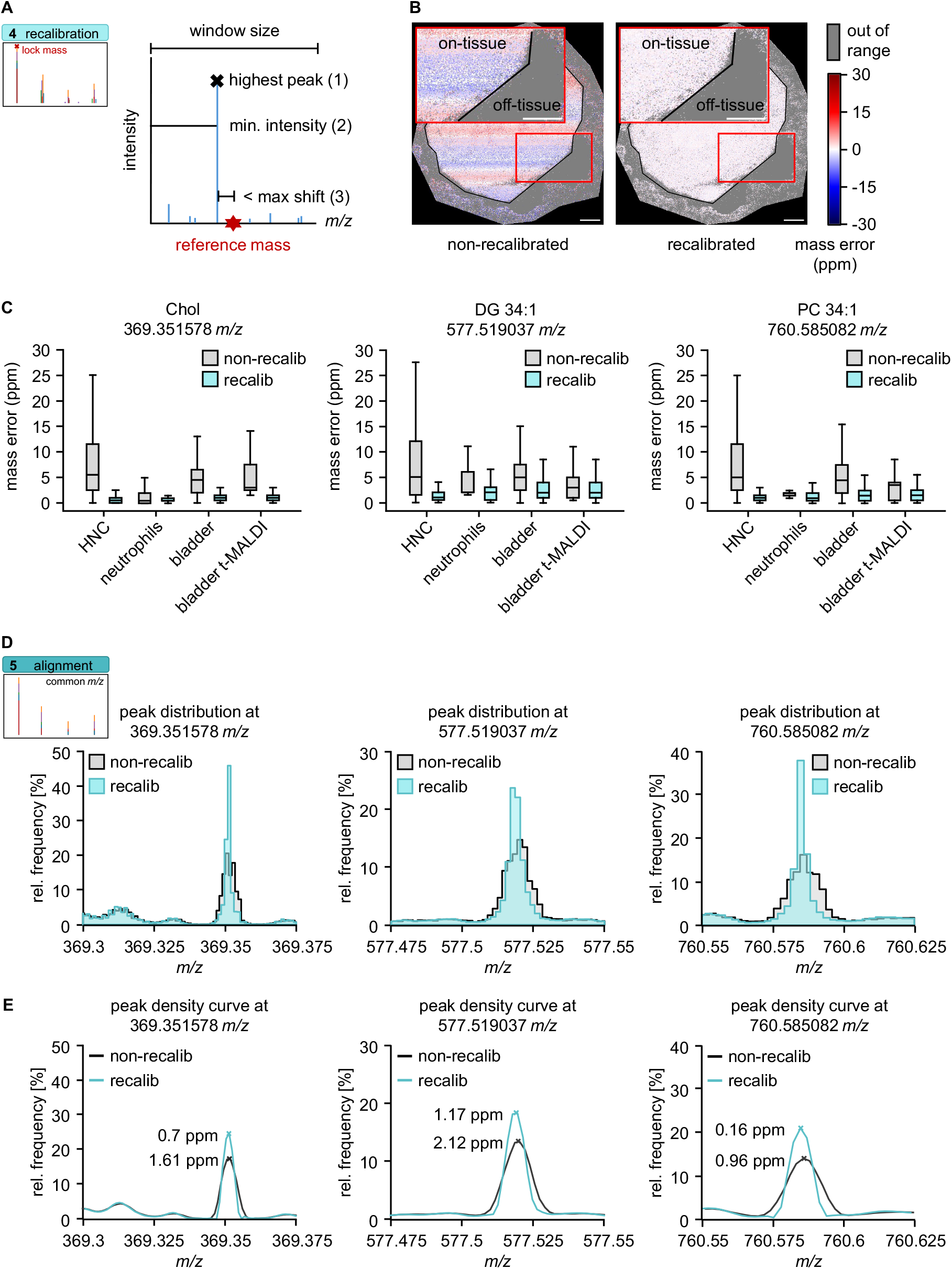
Automated mass recalibration in msiFlow 2.0 improves mass accuracy of pixel spectra and the common *m/z* vector. (**A**) Visualisation of automated peak selection for lock-mass-based recalibration. Experimental peaks are matched to predefined reference masses (red star) based on three selection criteria: (1) the highest peak intensity within a defined mass window, (2) a minimum intensity threshold, and (3) a maximum allowable mass shift from the reference mass. The mass differences between selected experimental peaks and their corresponding reference masses are used to calculate a recalibration function (median correction or linear regression), which is subsequently applied to correct the measured *m/z* values of the respective pixel spectrum. (**B**) Spatial distribution of pixel-wise mass error relative to *m/z* 369.351578 before (left) and after (right) recalibration in a representative bladder sample. Grey areas indicate pixels where no suitable peak was identified for recalibration according to the selection criteria shown in (A) and for which recalibration was therefore not performed. (**C**) Box plots showing the distribution of mass errors for all three reference masses (*m/z* 369.351578, 577.519037, and 760.585082) across successfully recalibrated pixels before (light grey) and after (light turquoise) recalibration (recalib) in four MSI datasets (HNC, *n* = 29; neutrophils, *n* = 6; bladder, *n* = 20; bladder t-MALDI, *n* = 2), highlighting the reduction in mass error achieved by recalibration. (**D**) Histograms showing the relative frequency distribution of detected peaks around the three reference masses before (light grey) and after (light turquoise) recalibration in the bladder MSI dataset. As a result of the recalibration, existing peaks are concentrated within a narrower *m/z* range around the reference masses, resulting in sharper peak-frequency maxima. (**E**) LOESS-smoothed peak-frequency curves derived from the histograms shown in (D). The maxima selected for construction of the common *m/z* vector are marked by crosses and labelled with their corresponding mass errors in ppm. Recalibration resulted in selected common *m/z* values with lower deviations from the respective reference masses. Scale bars: 300 µm.

We applied the recalibration workflow to four MALDI-2 and t-MALDI-2 MSI datasets using three reference masses distributed across the analysed *m/z* range. Recalibration reduced the pixel-wise mass errors of the three reference ions to approximately 0 ppm. A representative spatial heatmap for *m/z* 369.351578 illustrates this correction across the on-tissue area of a bladder section (Figure 2B) and the corresponding mass-error distributions confirm a consistent reduction across all four datasets (Figure 2C). Peaks fulfilling the matching criteria were detected in most on-tissue pixels, whereas off-tissue pixels generally lacked suitable reference peaks and were therefore not recalibrated. A maximum matching tolerance of 30 ppm was used based on the mass deviations observed during inspection of the raw data, as described in the Methods.

Because the reference masses were used to derive the recalibration function, their reduced mass errors primarily provide a technical verification that the recalibration procedure operated as intended. We therefore next examined whether recalibration also improved the subsequent construction of the common *m/z* vector during alignment. While recalibration corrects pixel-specific systematic mass deviations relative to known reference masses, alignment subsequently maps corresponding peaks across pixel spectra onto a common, data-derived *m/z* vector. Before recalibration, pixel-specific mass deviations produced relatively broad peak-frequency distributions around the reference masses. Following recalibration, the corresponding peaks were concentrated within narrower *m/z* intervals, resulting in sharper maxima in the peak-frequency distributions (Figure 2D). Consequently, the maxima of the density curves selected to construct the common *m/z* vector showed lower mass errors from the corresponding reference masses after recalibration (Figure 2E). Together, these results demonstrate that pixel-wise recalibration corrects systematic mass deviations at the reference masses and facilitates more precise construction of the common *m/z* vector. Successful recalibration nevertheless depends on the consistent detection of suitable reference ions with sufficient signal intensity and an appropriate distribution across the analysed mass range. Here, we used three reference masses that proved suitable for the analysed datasets measured by MALDI-2 in positive ion mode. However, their signal intensities may vary across sample types and may not be suitable for MALDI measurements without postionisation. Therefore, the suitability of these reference masses should be evaluated for each dataset. The spatial and statistical quality-control outputs generated by msiFlow 2.0 enable users to assess reference-ion coverage, identify spectra that cannot be recalibrated, and evaluate the resulting mass-error distributions. The effect of recalibration on independent MS/MS-supported lipid features was assessed in the subsequent performance evaluation.

### Performance Evaluation of msiFlow 2.0

We evaluated the performance of the updated alignment, pixel-wise mass recalibration, integrated lipid annotation, batch correction, and the computational optimisations implemented in msiFlow 2.0 using four MALDI-2 and t-MALDI-2 MSI datasets. MS/MS-supported lipid annotations available for the neutrophil and bladder datasets were used to independently assess mass accuracy and accurate-mass-based lipid annotation. For the mass-accuracy comparison, we included only lipid features detected by all three evaluated workflows: msiFlow 1.0, msiFlow 2.0 with updated alignment, and msiFlow 2.0 with recalibration followed by updated alignment. This resulted in 83 MS/MS-supported lipid features in the neutrophil dataset and 19 in the bladder dataset. The updated alignment significantly reduced absolute mass errors relative to msiFlow 1.0 in both datasets, with a further reduction observed after the addition of pixel-wise recalibration (Figure 3A). The combined recalibration and alignment workflow achieved mean absolute mass errors of 1.41 ppm in the neutrophil dataset and 1.13 ppm in the bladder dataset. These values show that the improvements observed for the reference masses in Figure 2 translated to independently assessed molecular features across a broader *m/z* range. For the neutrophil dataset, the mean absolute mass error of 1.41 ppm was within the low-ppm range previously reported for the same dataset following vendor-specific online lock-mass calibration during acquisition^29^. However, because the two approaches were not compared directly under identical processing conditions, these results should not be interpreted as a formal head-to-head comparison. Rather, they demonstrate that the open-source msiFlow 2.0 workflow achieves low-ppm mass accuracy while providing integrated visual quality-control outputs and direct processing of raw MSI data.

**Figure 3.**
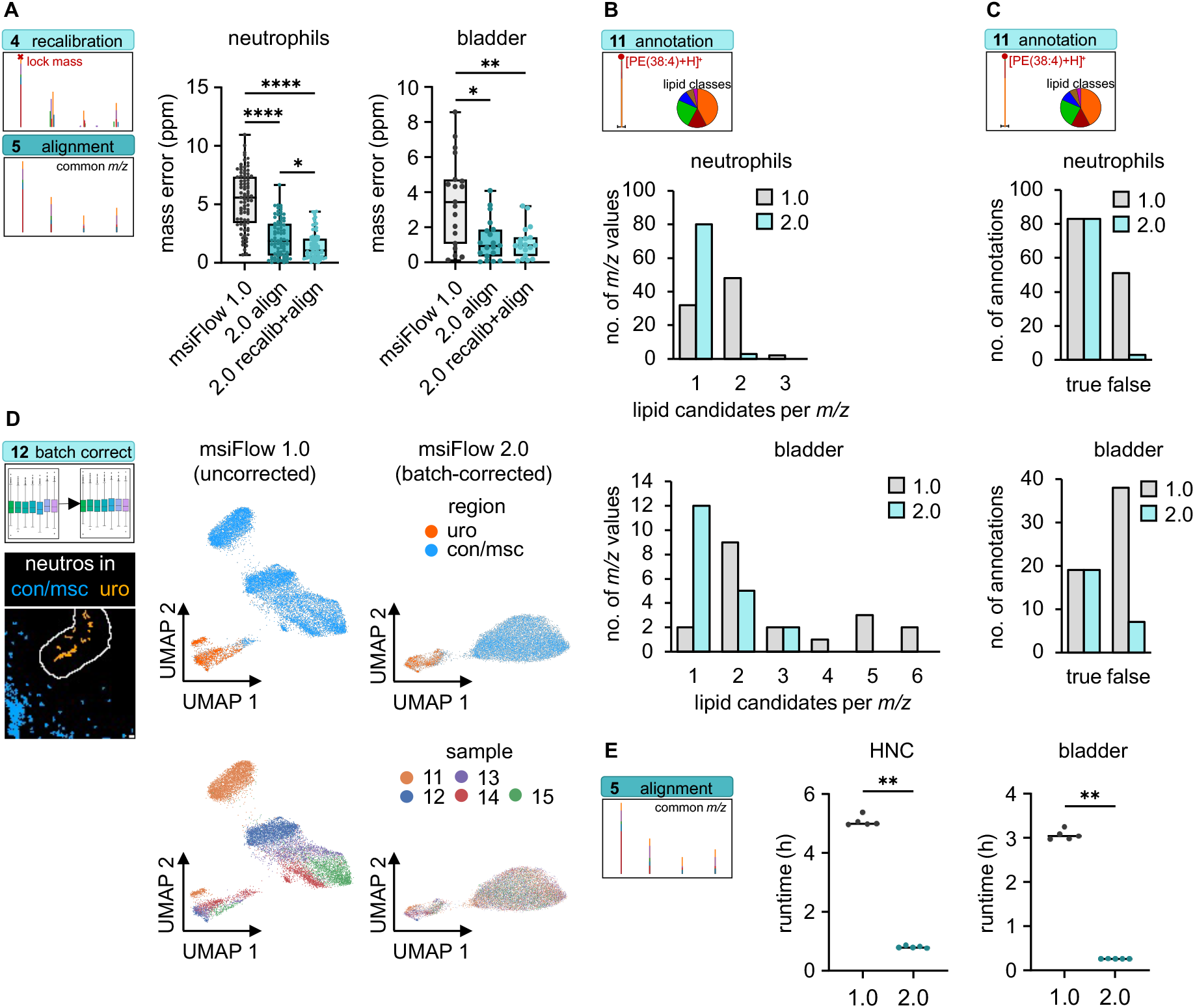
Performance evaluation of mass accuracy, lipid annotation, batch correction and computational efficiency in msiFlow 2.0. (**A**) Comparison of absolute mass errors obtained using msiFlow 1.0 (original workflow), msiFlow 2.0 align (updated alignment), and msiFlow 2.0 recalib+align (pixel-wise recalibration followed by updated alignment). The analysis included MS/MS-supported lipid features detected by all three workflows in the neutrophil (*n* = 83 MS/MS validated features) and bladder (*n* = 19 MS/MS validated features) MSI datasets. Box plots show the distributions of absolute mass errors for the three workflows. (**B**) Number of accurate-mass-based candidate annotations assigned to the MS/MS-supported lipid features in the neutrophil (upper) and bladder (lower) MSI datasets. The bar plot compares lipid candidate assignments obtained using the external annotation approach of msiFlow 1.0 (light grey) and the integrated msiFlow 2.0 annotation module (light turquoise). msiFlow 2.0 reduced the number of candidate annotations and increased the proportion of features with a unique candidate assignment. (**C**) Classification of candidate annotations according to their agreement with the corresponding MS/MS-supported lipid assignment in the neutrophil (upper) and bladder (lower) datasets. Compared with the msiFlow 1.0 annotation approach (light grey), msiFlow 2.0 (light turquoise) reduced the number of unsupported candidate annotations (false). (**D**) Effect of batch correction on the analysis of neutrophil pixels extracted from the bladder MSI dataset. Neutrophil pixels from a bladder MSI dataset were assigned to connective and muscle tissue (con/msc; blue) and urothelium (uro; orange) as shown by their spatial distributions. The corresponding UMAP embeddings of neutrophil pixels without batch correction (msiFlow 1.0, left) and with batch correction (msiFlow 2.0, right) show reduced sample-associated variation while retaining compartment-specific separation of the tissue compartments. Each point represents one neutrophil pixel. (**E**) Alignment runtime of the HNC and bladder datasets using msiFlow 1.0 and 2.0 in chunked processing mode. Each workflow was executed five times under identical computational conditions, revealing significant runtime reduction with msiFlow 2.0. Statistical analysis was performed as described in the Statistical Analysis section. * p<0.05, ** p<0.01, **** p<0.0001.

Next, we assessed how the integrated annotation module of msiFlow 2.0 affected the number of lipid candidates per *m/z*. In the original workflow, monoisotopic peak lists generated by msiFlow 1.0 were annotated externally using the LIPID MAPS bulk structure search. Although this search can be restricted by lipid class, adduct type, and mass tolerance, the available tolerance settings are based on fixed *m/z* increments. In contrast, msiFlow 2.0 supports ppm-based mass matching and additional carbon-number filtering within the automated workflow. Applying these criteria reduced the number of candidate annotations per *m/z* feature and increased the proportion of features with a unique candidate assignment in both the neutrophil and bladder datasets (Figure 3B).

To assess whether this reduction was consistent with the available experimental evidence, candidate annotations were compared with the corresponding MS/MS-supported lipid assignments. msiFlow 2.0 retained candidates consistent with the MS/MS-supported assignments while reducing the number of additional unsupported candidates (Figure 3C). Most excluded candidates were removed by restricting the search to lipid species with an even total carbon number, while further candidates were excluded by ppm-based mass matching (Tables S2 and S3). Notably, the reduction was more pronounced in the bladder dataset, for which three adduct types (M+H, M+Na, M+K) were considered, than in the neutrophil dataset, for which only (M+H) was included. This difference is expected because consideration of multiple adduct types increases the number of theoretical database matches for each measured *m/z*. Thus, the integrated annotation workflow reduces the candidate search space and facilitates more specific assignments. Nevertheless, accurate-mass matching alone cannot distinguish all isobaric or isomeric lipid structures, and the resulting assignments should therefore be interpreted as candidate annotations unless supported by orthogonal evidence such as MS/MS.

The effect of ComBat batch correction integrated in msiFlow 2.0 was examined using neutrophil pixels extracted from the bladder MSI dataset previously used to analyse the spatial neutrophil lipidome during UPEC infection^26^. As part of this analysis, neutrophil pixels in the bladder dataset were assigned to urothelium or connective and muscle tissue. Without batch correction, UMAP revealed two major compartment-associated clusters, while pixels from one sample separated from both principal clusters (Figure 3D).

Following ComBat correction, these pixels showed greater integration with the corresponding compartment-associated clusters, while the overall separation between urothelial and connective/muscle-tissue neutrophils was retained. These findings suggest that ComBat correction reduced sample-associated variation while retaining the major compartment-associated structure in the UMAP representation. This observation is consistent with recent evaluations identifying ComBat as a readily applicable batch-correction method for MALDI MSI^20, 21^. However, because the present evaluation is based primarily on the resulting low-dimensional embedding, it does not establish that all biological variation was preserved.

Finally, we assessed the computational performance of the updated alignment workflow using the HNC and bladder datasets, the two largest datasets evaluated, comprising 29 and 20 samples, respectively. Alignment was performed in five replicate computational runs for each workflow under identical hardware and resource settings. Compared with msiFlow 1.0, the updated workflow reduced alignment runtime by approximately 80% (Figure 3E). This corresponds to an approximately fivefold speed-up. In the original non-chunked implementation, construction of the global peak-frequency distribution required the complete *m/z* array to be held in memory. The corresponding memory requirements for the non-chunked processing mode were 563.8 GB for the HNC dataset and 228.3 GB for the bladder dataset, compared with 199.6 GB and 78.5 GB respectively, when using chunked processing of the updated workflow (Figure S5). By processing these arrays in smaller partitions in parallel, msiFlow 2.0 reduces the memory bottleneck and facilitates alignment of large MSI datasets. The magnitude of the performance improvement nevertheless depends on dataset size, storage performance, available memory, and the number of CPU cores.

Together, these results demonstrate that msiFlow 2.0 improves mass accuracy, reduces the number of unsupported candidate lipid annotations, mitigates sample-associated variation in downstream segmentation, and substantially improves the computational performance of alignment. The integration of these functions with visual quality control provides a more accurate, transparent, and scalable open-source framework for MSI data processing and analysis.

## Conclusions

We present msiFlow 2.0, a major release of the original msiFlow framework that integrates pixel-wise mass recalibration, automated lipid annotation, and batch correction, together with improved alignment, spatially informed deisotoping, computational scalability, and visual quality control. Evaluation using MALDI-2 and t-MALDI-2 MSI datasets demonstrated improved mass accuracy, fewer unsupported candidate lipid annotations, reduced sample-associated variation while retaining tissue compartment-associated variation in downstream segmentation, and substantially faster alignment of large datasets. By combining these functions within a configuration-driven and reproducible workflow, msiFlow 2.0 facilitates accurate and scalable MSI data processing while enabling transparent quality assessment at multiple stages of the analysis. The open-source implementation provides an accessible framework for the analysis and biological interpretation of spatial lipidomic data across different MSI settings.

## Associated Content

### Data Availability Statement

All MALDI MSI datasets used in this study are publicly available. The previously published neutrophil data and bladder data of control and UPEC-infected mice are available through the Zenodo repositories and accession numbers provided in the corresponding publications. The remaining MALDI MSI data of mouse bladders under TGF-β1 inhibition and the HNC data are deposited in the research data repository of the University of Münster (datastore) and are available at https://doi.org/10.17879/b2892-0g370 and https://doi.org/10.17879/t3vmd-s6708. The t-MALDI MSI bladder data is also available on datastore at https://doi.org/10.17879/40rcy-va712.

### Code Availability Statement

The source code of msiFlow 2.0 used in this study is publicly available as a release on GitHub (https://github.com/Immunodynamics-Engel-Lab/msiflow) and permanently archived in Zenodo (https://doi.org/10.5281/zenodo.23923290). A versioned Docker image containing the complete software environment is available on Docker Hub: phispa1812/msiflow_cli:v2.0.0 (CLI version) and phispa1812/msiflow_gui:v2.0.0 (GUI version).

### Supporting Information

Supporting Information includes quality-control outputs, memory-usage comparison, HNC clinicopathological characteristics, and lipid annotation tables.

### Author Information

#### Author Contributions

Conceptualisation: PS, DRE

Methodology: PS, S. Bessler, JS, DRE

Software: PS

Validation: PS, DRE Formal analysis: PS

Investigation: PS, S. Bessler, EMH, LW, JB, ST, AP

Resources: S. Bessler, EMH, AP, S. Brandau, JS

Data curation: PS, S. Bessler, EMH

Writing – original draft: PS

Writing – review & editing: all authors

Visualisation: PS

Supervision: DRE

Project administration: DRE

Funding acquisition: PS, S. Brandau, DRE

### Funding

We received funding from the German Research Foundation: FOR5427 (466687329) SP4 (DRE); EN984/16-1, 18-1 and 19-1 (DRE); TR332 (449437943) A3 (DRE), Z1 (OS) and A4 (SBr); INST 20876/486-1 (DRE), SFB1752 (549898653) B05 (DRE), Graute-Oppermann-Graduierten Kolleg (PhaSe) (DRE) and the BIOME Postdoctoral Excellence Programme (BIOME PEP) of the Faculty of Medicine, University of Duisburg-Essen (PS).

### Notes

The authors declare no competing financial interest.

## Acknowledgments

We acknowledge the excellent technical assistance of Mathias Schmidt and support by the Open Access Publication Fund of the University of Duisburg-Essen, the Central Animal Facilities of the Medical Faculty Essen, the Imaging Center Essen (IMCES), the Immunoproteomics group and the mass spectrometry unit MOSAIC. Mass spectrometry measurements were supported by Bruker Daltonics. Calculations for this publication were performed on HPC cluster PALMA II of the University of Münster, subsidised by the DFG (INST 211/667-1).

## References

(1) Soltwisch, J.; Kettling, H.; Vens-Cappell, S.; Wiegelmann, M.; Müthing, J.; Dreisewerd, K. Mass spectrometry imaging with laser-induced postionization. Science 2015, 348 (6231), 211–215.

(2) Niehaus, M.; Soltwisch, J.; Belov, M.; Dreisewerd, K. Transmission-mode MALDI-2 mass spectrometry imaging of cells and tissues at subcellular resolution. Nat. Methods 2019, 16 (9), 925–931.

(3) Dreisewerd, K.; Bien, T.; Soltwisch, J. MALDI-2 and t-MALDI-2 mass spectrometry imaging. In *Mass Spectrometry Imaging of Small Molecules: Methods and Protocols*, Springer, 2021; pp 21–40.

(4) Balluff, B.; Hopf, C.; Porta Siegel, T.; Grabsch, H. I.; Heeren, R. M. Batch effects in MALDI mass spectrometry imaging. J. Am. Soc. Mass Spectrom. 2021, 32 (3), 628–635.

(5) Weiskirchen, R.; Weiskirchen, S.; Kim, P.; Winkler, R. Software solutions for evaluation and visualization of laser ablation inductively coupled plasma mass spectrometry imaging (LA-ICP-MSI) data: a short overview. J. Cheminform. 2019, 11 (1), 16.

(6) Hu, H.; Laskin, J. Emerging computational methods in mass spectrometry imaging. Adv. Sci. 2022, 9 (34), 2203339.

(7) Veselkov, K.; Sleeman, J.; Claude, E.; Vissers, J. P. C.; Galea, D.; Mroz, A.; Laponogov, I.; Towers, M.; Tonge, R.; Mirnezami, R.;, et al. BASIS: High-performance bioinformatics platform for processing of large-scale mass spectrometry imaging data in chemically augmented histology. Sci. Rep. 2018, 8 (1), 4053. DOI: 10.1038/s41598-018-22499-z.

(8) Cordes, J.; Enzlein, T.; Marsching, C.; Hinze, M.; Engelhardt, S.; Hopf, C.; Wolf, I. M2aia—Interactive, fast, and memory-efficient analysis of 2D and 3D multi-modal mass spectrometry imaging data. GigaScience 2021, 10 (7), giab049.

(9) Guo, G.; Papanicolaou, M.; Demarais, N.; Wang, Z.; Schey, K.; Timpson, P.; Cox, T.; Grey, A. Automated annotation and visualisation of high-resolution spatial proteomic mass spectrometry imaging data using HIT-MAP. Nat. Commun. 2021, 12 (1), 3241.

(10) Bond, N. J.; Koulman, A.; Griffin, J. L.; Hall, Z. massPix: an R package for annotation and interpretation of mass spectrometry imaging data for lipidomics. Metabolomics 2017, 13, 1–5.

(11) Robichaud, G.; Garrard, K. P.; Barry, J. A.; Muddiman, D. C. MSiReader: an open-source interface to view and analyze high resolving power MS imaging files on Matlab platform. J. Am. Soc. Mass Spectrom. 2013, 24 (5), 718–721. DOI: 10.1007/s13361-013-0607-z From NLM Medline.

(12) Race, A. M.; Palmer, A. D.; Dexter, A.; Steven, R. T.; Styles, I. B.; Bunch, J. SpectralAnalysis: software for the masses. Anal. Chem. 2016, 88 (19), 9451–9458.

(13) Bemis, K. D.; Harry, A.; Eberlin, L. S.; Ferreira, C.; van de Ven, S. M.; Mallick, P.; Stolowitz, M.; Vitek, O. Cardinal: an R package for statistical analysis of mass spectrometry-based imaging experiments. Bioinformatics 2015, 31 (14), 2418–2420.

(14) Bemis, K. A.; Föll, M. C.; Guo, D.; Lakkimsetty, S. S.; Vitek, O. Cardinal v. 3: a versatile open-source software for mass spectrometry imaging analysis. Nat. Methods 2023, 20 (12), 1883–1886.

(15) Föll, M. C.; Moritz, L.; Wollmann, T.; Stillger, M. N.; Vockert, N.; Werner, M.; Bronsert, P.; Rohr, K.; Grüning, B. A.; Schilling, O. Accessible and reproducible mass spectrometry imaging data analysis in Galaxy. GigaScience 2019, 8 (12). DOI: 10.1093/gigascience/giz143.

(16) Galaxy for accessible, reproducible, and collaborative data analyses: 2026 update. Nucleic Acids Res. 2026, gkag469.

(17) Köster, J.; Rahmann, S. Snakemake—a scalable bioinformatics workflow engine. Bioinformatics 2012, 28 (19), 2520–2522.

(18) Fonville, J. M.; Carter, C.; Cloarec, O.; Nicholson, J. K.; Lindon, J. C.; Bunch, J.; Holmes, E. Robust data processing and normalization strategy for MALDI mass spectrometric imaging. Anal. Chem. 2012, 84 (3), 1310–1319.

(19) Johnson, W. E.; Li, C.; Rabinovic, A. Adjusting batch effects in microarray expression data using empirical Bayes methods. Biostatistics 2007, 8 (1), 118–127. DOI: 10.1093/biostatistics/kxj037.

(20) Metodiev, M.; Dexter, A.; Zhou, W.; González-Fernández, A.; Nikula, C.; Johns, L. M.; Karali, E.; Kazanc, E.; Tsalikis, A.; Tripp, A. Evaluating batch correction methods for large-scale mass spectrometry imaging of heterogeneous tissues. Anal. Chem. 2026, 98 (5), 3531.

(21) Sparre, A. A.; Jensen, O. N. Batch correction for large-scale mass spectrometry imaging experiments. Bioinformatics 2026, 42 (6). DOI: 10.1093/bioinformatics/btag360.

(22) Treu, A.; Römpp, A. Matrix ions as internal standard for high mass accuracy matrix-assisted laser desorption/ionization mass spectrometry imaging. Rapid Commun. Mass Spectrom. 2021, 35 (16), e9110.

(23) Inglese, P.; Huang, H. X.; Wu, V.; Lewis, M. R.; Takats, Z. Mass recalibration for desorption electrospray ionization mass spectrometry imaging using endogenous reference ions. BMC Bioinformatics 2022, 23 (1), 133.

(24) Spangenberg, P.; Bessler, S.; Widera, L.; Bottek, J.; Richter, M.; Thiebes, S.; Siemes, D.; Krauß, S. D.; Migas, L. G.; Kasarla, S. S. msiFlow: automated workflows for reproducible and scalable multimodal mass spectrometry imaging and microscopy data analysis. Nat. Commun. 2025, 16 (1), 1065.

(25) Merkel, D. Docker: lightweight linux containers for consistent development and deployment. Linux journal 2014, 2014 (239), 2.

(26) Widera, L.; Spangenberg, P.; Siemes, D.; Bessler, S.; Bottek, J.; Voss, H.; Novak, A.; Hueser, D.; Richter, M.; Potthoff, A. TSP1/TGF-β1 drives arachidonic acid metabolism to orchestrate neutrophil swarming. bioRxiv 2025, 2025.2011. 2025.688397.

(27) Truong, J. X.; Spotbeen, X.; White, J.; Swinnen, J. V.; Butler, L. M.; Snel, M. F.; Trim, P. J. Removal of optimal cutting temperature (OCT) compound from embedded tissue for MALDI imaging of lipids: Truong JXM et al. Anal. Bioanal. Chem. 2021, 413 (10), 2695–2708.

(28) Potthoff, A.; Schwenzfeier, J.; Niehaus, M.; Bessler, S.; Hoffmann, E.; Soehnlein, O.; Höhndorf, J.; Dreisewerd, K.; Soltwisch, J. Spatial biology using single-cell mass spectrometry imaging and integrated microscopy. Nat. Commun. 2025, 16 (1), 9129.

(29) Bessler, S.; Richter, M.; Schwenzfeier, J.; Reinartz Groba, S. N.; Trebicka, J.; Dreisewerd, K.; Soltwisch, J.; Soehnlein, O. Integrated Single-Cell Mass Spectrometry Imaging and Immunofluorescence Microscopy for Multimodal Characterization of Human Immune Cells. Anal. Chem. 2026, 98 (14), 10338–10344. DOI: 10.1021/acs.analchem.6c00381.

(30) Behdenna, A.; Colange, M.; Haziza, J.; Gema, A.; Appé, G.; Azencott, C.-A.; Nordor, A. pyComBat, a Python tool for batch effects correction in high-throughput molecular data using empirical Bayes methods. BMC Bioinformatics 2023, 24 (1), 459.

(31) Leland, M.; John, H.; Nathaniel, S.; Lukas, G. UMAP: uniform manifold approximation and projection. J. Open Source Softw. 2018, 3 (29), 861.

(32) McInnes, L.; Healy, J.; Astels, S. hdbscan: Hierarchical density based clustering. J. Open Source Softw. 2017, 2 (11), 205.

(33) Palmer, A.; Phapale, P.; Chernyavsky, I.; Lavigne, R.; Fay, D.; Tarasov, A.; Kovalev, V.; Fuchser, J.; Nikolenko, S.; Pineau, C. FDR-controlled metabolite annotation for high-resolution imaging mass spectrometry. Nat. Methods 2017, 14 (1), 57–60.

(34) Strohalm, M.; Kavan, D.; Novák, P.; Volný, M.; Havlíček, V. r. mMass 3: A Cross-Platform Software Environment for Precise Analysis of Mass Spectrometric Data. Anal. Chem. 2010, 82 (11), 4648–4651. DOI: 10.1021/ac100818g.

