## Supporting Information for "msiFlow 2.0: Extending Mass Spectrometry Imaging Workflows with Mass Recalibration, Annotation, Batch Correction, and Visual Quality Control"


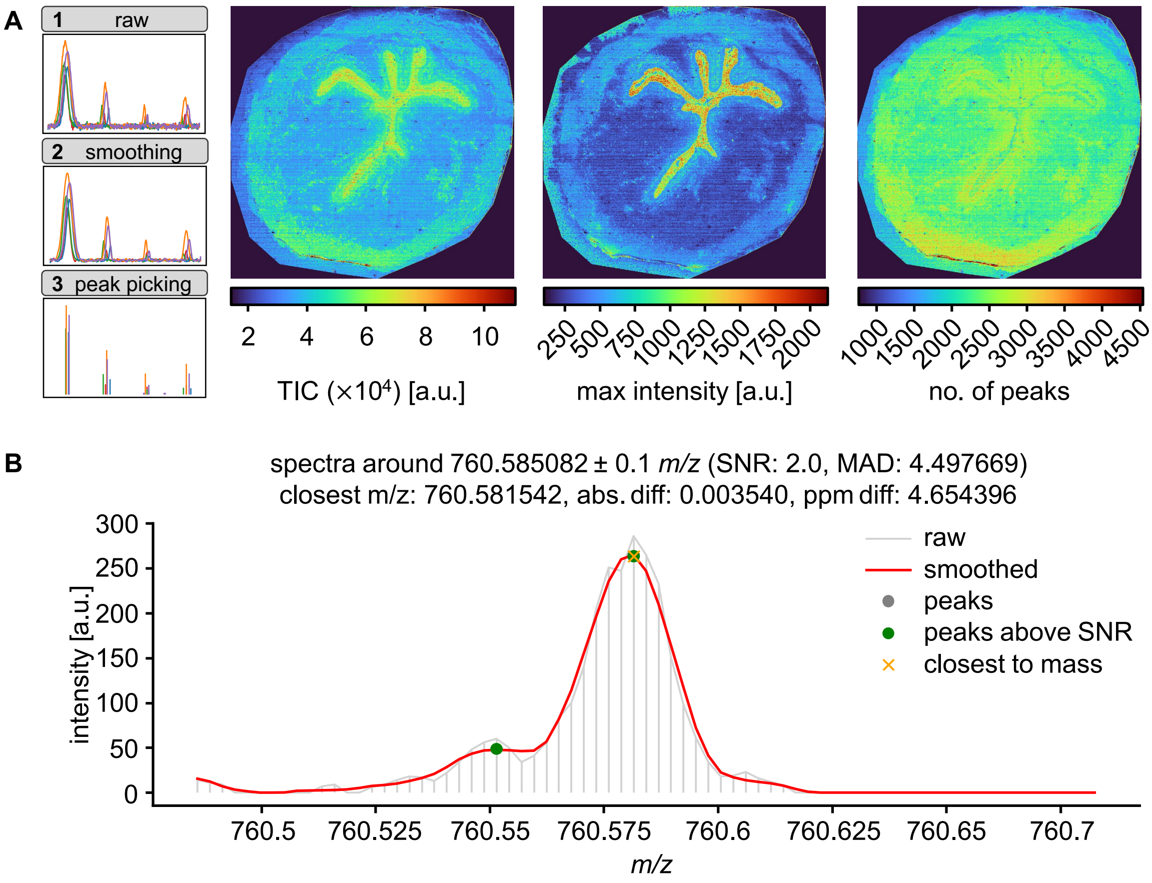


**Figure S1. Visual quality-control outputs for raw MSI data, spectral smoothing, and peak picking in msiFlow 2.0.** (**A**) Spatial heatmaps showing total ion current (left), maximum peak intensity (centre) and number of detected peaks (right) per pixel in a representative bladder sample measured by MALDI-2 MSI. (**B**) Representative raw (grey) and smoothed (red) spectra within a user-defined mass window centred at *m/z* 760.585082 ± 0.1 *m/z*. Peaks detected in the smoothed spectra (green). The detected peak closest to the user-defined *m/z* value is highlighted (orange x). TIC, total ion current; SNR, signal-to-noise ratio; MAD, median absolute deviation; abs. diff., absolute mass difference; ppm diff., mass difference in ppm.

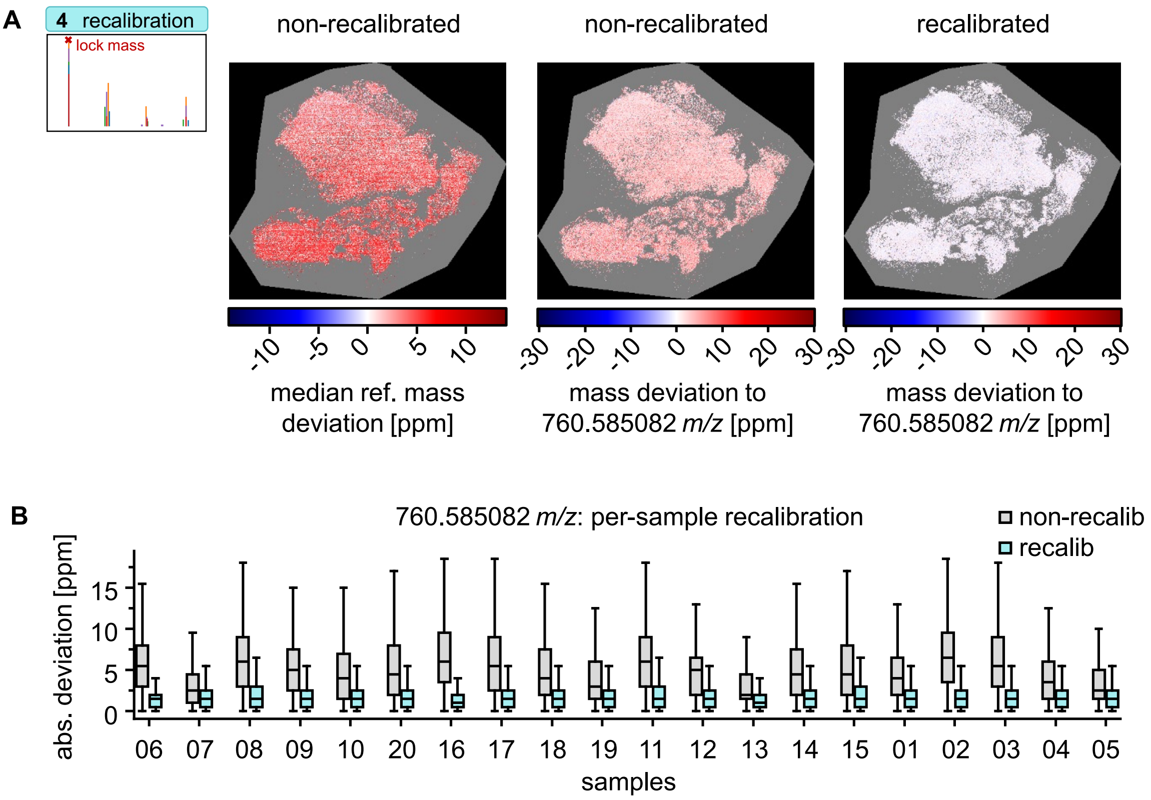


**Figure S2. Visual quality control of pixel-wise mass recalibration in msiFlow 2.0.** (**A**) Spatial heatmaps showing the median mass deviation across all reference masses (left) and the mass error relative to the reference mass at *m/z* 760.585082 before (centre) and after (right) recalibration in a representative head and neck cancer sample acquired by MALDI-2 MSI. (**B**) Box plots showing the pixel-wise distributions of the absolute mass deviation to *m/z* 760.585082 before and after recalibration for each sample in the bladder dataset ($n$ = 20). ref, reference; abs, absolute; recalib, recalibrated.

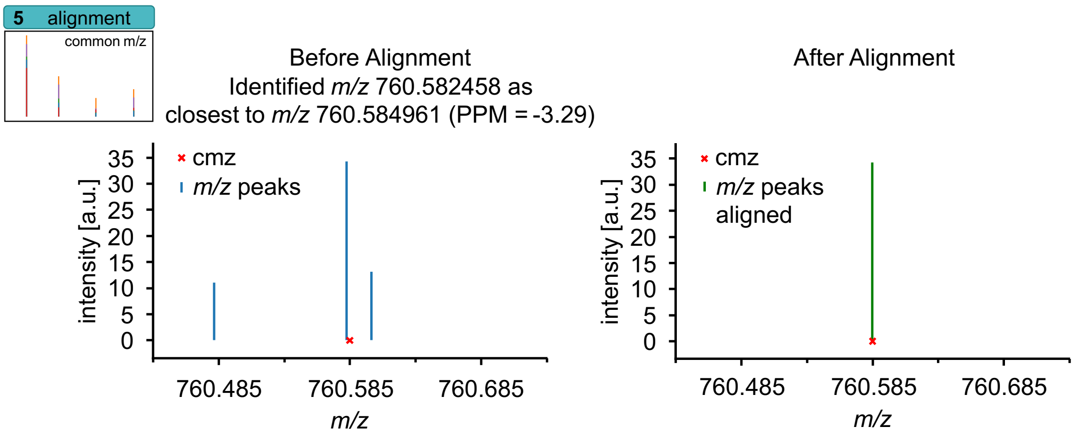


**Figure S3. Visual quality-control outputs for alignment in msiFlow 2.0.** Additional output related to Figure 2D, E. Peak-picked data of one selected pixel with a certain *m/z* range. The *m/z* peaks are shown before (left, blue peaks) and after (right, green peak) alignment at the selected mass (*m/z* 760.584961) in relation to the reference mass *m/z* 760.585082. Only the peak at *m/z* 760.582458 (left, highest blue peak), which is closest to the selected common-axis value at *m/z* 760.584961 (red cross), is mapped to this value. The remaining peaks are not mapped because they fall outside the specified matching tolerance.

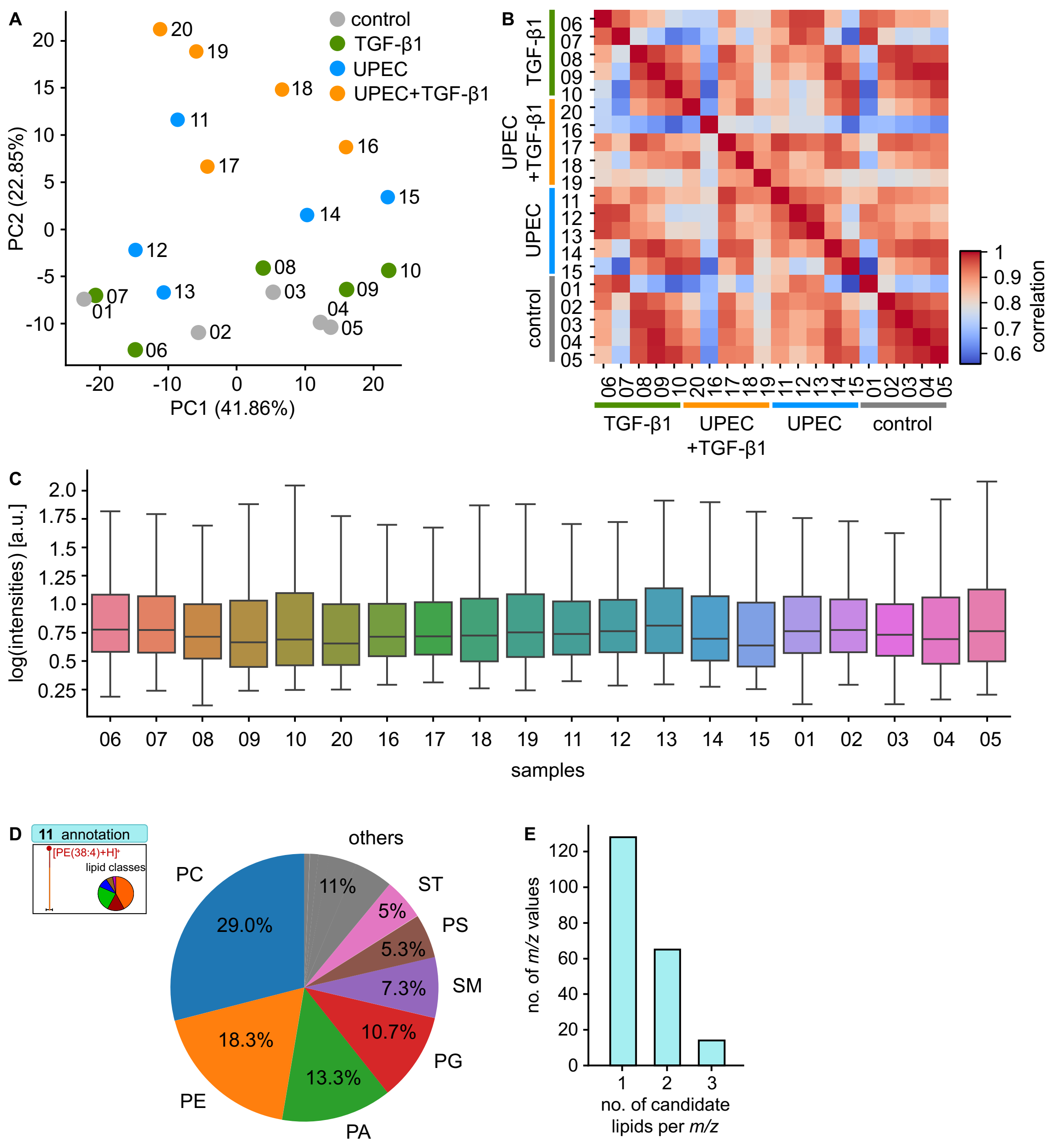


**Figure S4. Summary plots and lipid-annotation outputs provided by msiFlow 2.0 shown for the bladder dataset.** Mean feature intensities were calculated across all pixels (on- and off-tissue) within each sample. The resulting sample-by-feature intensity matrix was used to generate (**A**) principal component analysis (PCA) plot using all features, (**B**) a heatmap showing the Pearson correlation between all feature intensities per sample, and (**C**) box plots showing the distribution of $\log_{10}$-transformed mean feature intensities for each sample. (**D**) Pie chart showing the distribution of all candidate lipid annotations across lipid classes, including multiple assignments associated with the same *m/z* feature. (**E**) Histogram showing the number of candidate lipid annotations assigned to each measured *m/z* value.

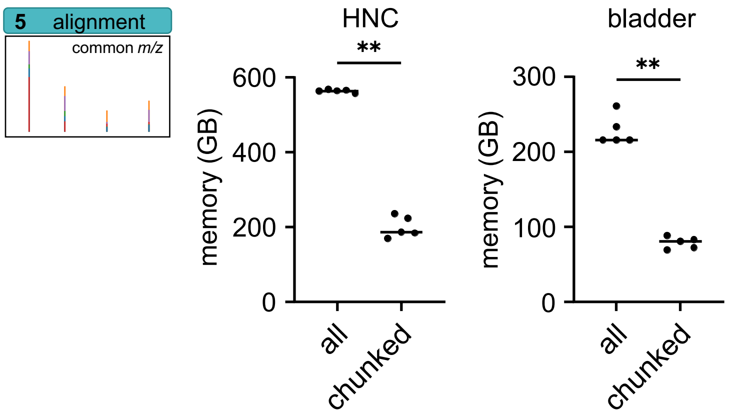


**Figure S5. Memory usage of the two processing modes of the alignment workflow in msiFlow 2.0.** Memory usage for the HNC (left) and bladder (right) datasets was assessed across five replicate computational runs under identical hardware and resource settings using the two processing modes: all, in which the entire dataset is loaded into memory, and chunked, in which the data are processed in smaller chunks. Statistical analysis was performed as described in the Statistical Analysis section. ** $p<0.01$.

**Table S1. Clinicopathological Characteristics of Patients with Head and Neck Cancer**

| | n | % |
|---|---|---|
| **All samples** | 29 | 100 |
| **Sex** | | |
| Male | 21 | 72.4 |
| Female | 8 | 27.6 |
| **Age, years** | | |
| Mean ± SD | 64.3 ± 8.6 | |
| Median (range) | 64 (49-87) | |
| **Tumour localisation** | | |
| Oropharynx | 21 | 72.4 |
| Oral cavity | 8 | 27.6 |
| **T category** | | |
| T1 | 3 | 10.3 |
| T2 | 9 | 31.0 |
| T3 | 7 | 24.1 |
| T4/T4a | 10 | 34.5 |
| **N category** | | |
| N0 | 9 | 31.0 |
| N1 | 5 | 17.2 |
| N2/N2a-c | 12 | 41.4 |
| N3/N3b | 3 | 10.3 |
| **Distant metastasis** | | |
| M0 | 28 | 96.6 |
| M1 | 1 | 3.4 |
| **UICC stage** | | |
| Stage I | 6 | 20.7 |
| Stage II | 6 | 20.7 |
| Stage III | 5 | 17.2 |
| Stage IV (not further specified) | 1 | 3.4 |
| Stage IVA | 9 | 31.0 |
| Stage IVB | 2 | 6.9 |
| **Histological grading** | | |
| G2 | 14 | 48.3 |
| G3 | 8 | 27.6 |
| Unknown/not reported | 7 | 24.1 |
| **p16/HPV status (as recorded)** | | |
| p16-positive | 13 | 44.8 |
| p16-negative | 15 | 51.7 |
| HPV-positive | 1 | 3.4 |

**Table S2. Lipid annotation examples from neutrophil dataset**

| | | msiFlow 1.0 | | | | msiFlow 2.0 | | | |
|---|---|---|---|---|---|---|---|---|---|
| lipid | theoretical *m/z* | *m/z* | mass error [ppm] | lipid candidate | adduct | *m/z* | mass error [ppm] | lipid candidate | adduct |
| PC O-40:6 | 820.621467 | 820.6157 | 7.03 | PS 38:0<br>PC O-40:6 | H<br>H | 820.6198 | 2.03 | PC O-40:6 | H |
| PC 40:5 | 836.616381 | 836.61 | 7.63 | PC 40:5<br>Hex2Cer 32:0;O2 | H<br>H | 836.61365 | 3.26 | PC 40:5 | H |
| Hex2Cer 34:1;O2 | 862.625018 | 862.6237 | 1.53 | PC 42:6<br>Hex2Cer 34:1;O2 | H<br>H | 862.625 | 0.02 | Hex2Cer 34:1;O2 | H |

**Table S3. Lipid annotation examples from bladder dataset**

| | | msiFlow 1.0 | | | | msiFlow 2.0 | | | |
|---|---|---|---|---|---|---|---|---|---|
| lipid | theoretical m/z | m/z | mass error [ppm] | lipid candidate | adduct | m/z | mass error [ppm] | lipid candidate | adduct |
| PC 32:1 | 732.553781 | 732.553 | 1.07 | PE 35:1<br>PC 32:1<br>HexCer 34:1;O4 | H<br>H<br>H | 732.5539 | 0.16 | PC 32:1 | H |
| PE 38:5 | 766.538131 | 766.54144 | 4.32 | PE O-38:6<br>PC 35:5<br>PE 38:5<br>HexCer 35:2<br>PC 33:2<br>PE 36:2 | H<br>H<br>H<br>Na<br>Na<br>Na | 766.5405 | 3.09 | PE O-38:6;O<br>PE 38:5 | H<br>H |
| PC 36:4 | 782.569431 | 782.56573 | 4.73 | PC 36:4<br>PE 39:4<br>PE 37:1<br>PC 34:1<br>HexCer 36:1 | H<br>H<br>Na<br>Na<br>Na | 782.56866 | 0.95 | PC 36:4<br>PC 34:1 | H<br>Na |
| PC 38:4 | 810.600731 | 810.5994 | 1.64 | PC 38:4<br>PE 41:4<br>PE 39:1<br>PC 36:1<br>HexCer 38:1;O4 | H<br>H<br>Na<br>Na<br>Na | 810.60004 | 0.85 | PC 38:4<br>PC 36:1 | H<br>Na |
